# The anisotropic sensitivities of perceptual speed between expanding and contracting optic flows

**DOI:** 10.64898/2026.01.13.699182

**Authors:** Junxiang Luo, Hiroshi Ashida, Isao Yokoi, Hiromasa Takemura

## Abstract

Retinal optic flow provides critical visual cues for locomotion and self-navigation. Forward self-motion, such as walking or driving, typically induces an expanding radial flow pattern on the retina. Despite the predominance of expanding optic flow in everyday self-motion, several studies have reported higher perceptual sensitivity for contracting flow that arises under relatively rare conditions, such as walking backward or falling onto one’s back. The underlying basis of this perceptual anisotropy remains elusive. In this study, we compared perceptual sensitivity to speed for expanding versus contracting optic flow, given that speed is a fundamental attribute of motion and that perceptual sensitivity to it provides insight into underlying visual mechanisms. We found that sensitivity in speed discrimination was comparable between expansion and contraction at low and high speeds, but was significantly higher for contraction at an intermediate speed corresponding to ecologically relevant self-motion, such as walking. This anisotropic pattern was observed for both 2D planar (Experiment 1) and simulated 3D (Experiment 2) optic flow stimuli. Together, these findings suggest that anisotropic perceptual sensitivity to expanding and contracting optic flow at ecologically typical egomotion speeds may reflect an adaptive visual mechanism that supports postural stability and balance during relatively rare backward self-motion.

## Introduction

Information about self-generated body movements is essential for supporting key behaviors of humans and other animals, including postural control, collision avoidance, and spatial navigation. Optic flow — the pattern of visual motion projected onto the retina during observer movement — serves as a critical visual cue for self-motion perception (Gibson, 2015). During forward movements such as walking or running, the retina receives an expanding optic flow pattern in which visual elements radiate outward from a central point. Conversely, backward movement, although less common in everyday life, produces a contracting pattern in which visual elements move inward toward a central point on the retina. These two radial patterns, termed expansion and contraction, engage shared neuronal substrates and visual processing mechanisms in motion-sensitive cortical areas (Duffy & Wurtz, 1991; Morrone et al., 2000; Saito et al., 1986; Smith, Wall, Williams, & Singh, 2006; Tanaka, Fukada, & Saito, 1989).

However, even though both types of radial optic flow recruit broadly similar neural mechanisms, a fundamental unresolved question is whether their processing also diverges at some stage, given their different frequencies of occurrence in natural behavior and their opposite motion trajectories. Psychophysical studies have compared perceptual sensitivities between expansion and contraction, yet their findings remain inconsistent. Some studies reported greater perceptual sensitivity to contraction (Edwards & Badcock, 1993; Edwards & Ibbotson, 2007; Geesaman & Qian, 1998), others found greater sensitivity to expansion (Clifford, Beardsley, & Vaina, 1999; Lopez-Moliner, 2005), and still others didn’t report reliable difference (Beardsley & Vaina, 2005; Morrone, Burr, Di Pietro, & Stefanelli, 1999). These discrepancies may arise from variations in stimulus parameters and also from methodological differences in how perceptual sensitivity is assessed.

Importantly, simply detecting or differentiating radial optic flow patterns is not sufficient for adaptive behavior. Self-movements such as walking, running, and driving generate optic flows with different speeds, and effective navigation requires the visual system to estimate not only the type but also the speed of the optic flow. Accurate estimation of optic flow speed underlies crucial behaviors such as regulating locomotion, estimating time-to-collision, and maintaining stability during rapid movement. Therefore, to better understand the mechanisms of self-motion perception, it is necessary to characterize how perceptual sensitivity to optic-flow speed varies across expansion and contraction patterns and across different speed ranges.

In the present study, we asked whether human observers exhibit distinct perceptual sensitivities to optic-flow speed across a broad range of speeds in expanding and contracting radial patterns. We conducted two psychophysical experiments in which participants performed speed discrimination tasks on radial optic flow stimuli spanning speeds typical of walking and driving, as well as speeds much slower and faster than those typically experienced. Perceptual sensitivity was quantified using the Weber fraction, which describes discrimination performance in accordance with Weber’s Law (Theodor, 1966). In Experiment 1, we used 2D planar radial motions with uniform speed across the visual field to strictly control local speed components at all eccentricities. In Experiment 2, we generated 3D-compatible optic flow fields that simulated more naturalistic self-motion. For each experiment, we estimated speed-discrimination sensitivity at multiple reference speeds for both expansion and contraction.

Across both experiments, we found greater perceptual sensitivity to contraction than to expansion at medium speeds, corresponding to typical everyday locomotion. This difference was not significant at very low and very high speeds. Such a pattern may reflect an adaptive characteristic of the visual system, potentially tuned to ecologically relevant self-motion conditions where contraction (e.g., during backward instability or falling) demands heightened vigilance.

## Experiment 1

In Experiment 1, we measured speed discrimination for 2D planar radial flow patterns that maintain a constant speed between dots and across the entire visual field.

### Methods

#### Participants

Twenty-seven healthy human adults (16 females and 11 males, aged 28.6 ± 1.4 S.E. years, ranging from 20 to 42 years old) participated in the experiment. All of them have normal or corrected-to-normal visual acuity. A written informed consent form, including instructions on experimental safety, voluntary participation, the right to withdraw, and agreement to make anonymized data available to the public, was obtained from all participants before the experiment. All participants underwent a test evaluating their visual acuity, astigmatism, and binocular stereopsis before the main experiments. The study protocol was approved by the Ethics Committee of the National Institutes of Natural Sciences (protocol number: EC01-64).

#### Apparatus

Visual stimuli were generated on a workstation (Dell Precision 3650, Dell, Round Rock, TX) running Linux (Ubuntu 22.04 LTS). To present visual stimuli, we used a PROPixx DLP LED projector (VPixx Technologies, Saint-Bruno, QC, Canada) with a refresh rate of 1440 Hz and a spatial resolution of 1920 × 1080 pixels. Screen luminance was measured using a colorimeter (CS-150, Konica Minolta Inc., Japan). The minimum (black level) and maximum (white level) luminances are approximately 4 and 8500 cd/m^2^, respectively. Presenting ultra-fast motion stimuli with the 1440 Hz projector minimized the apparent-motion-type artificial effect, bringing the sensation of ultra-fast optic flow closer to natural conditions. Visual stimuli were projected onto a screen, forming a rectangular display measuring 37.5 cm wide by 21 cm high. A chinrest was mounted at the front of the projection screen. Participants were asked to rest their heads on it to stabilize their head position and maintain a constant observation distance. The distance between the eyes and the projection screen is 93 cm, resulting in a visual angle of approximately 22.80° (width) × 12.88° (height). All experiments were conducted in a dark room enclosed by light-blocking curtains to minimize ambient light. Participants reported their perceptions by pressing the L or R button on a Retro-Bit Legacy 16 gaming pad (Retro-Bit, Pomona, CA). Fixation was monitored using an eye-tracking system (LiveTrack Lightning, Cambridge Research Systems, Rochester, UK). The eye-tracking camera was positioned 18 cm in front of the participant’s eyes, aligned with the chinrest. Binocular eye positions were recorded online at a sampling rate of 500 Hz.

#### Visual stimuli

Visual stimuli were designed using MATLAB Psychtoolbox extensions (Brainard, 1997; Kleiner, Brainard, & Pelli, 2007; http://psychtoolbox.org/). **Figure 1A** describes a schematic illustration of the stimuli used in the experiment. The stimuli consisted of a two-dimensional random dot field, drawn in white and presented against a black background. Each dot is 0.2° in size, corresponding to around 17 pixels on the screen. The density of the dot field is 0.3 dots per square degree. Dots were initially placed at random positions on the screen and then moved radially. Two stimulus conditions were tested: in the ‘expansion’ condition, dots moved outward from the center (**Supplementary Video 1**); in the ‘contraction’ condition, they moved inward toward the center (**Supplementary Video 2**). Each dot traveled a distance of 5° before being removed and reinitialized to a new random location. All dots moved at a uniform speed across the visual field. During stimulus presentation, a white fixation cross of 0.5° in diameter was presented at the center of the screen.

**Figure 1.**
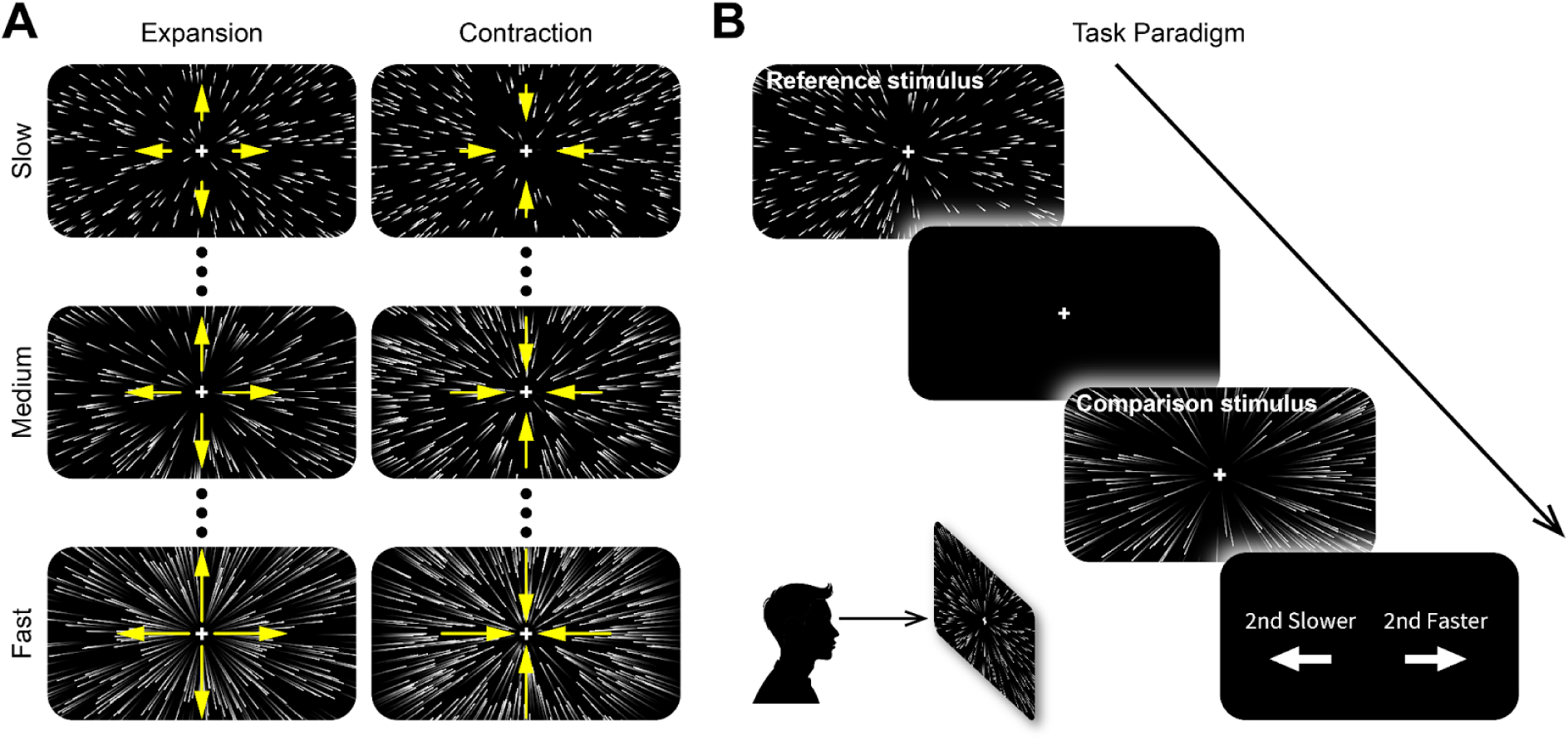
Schematic illustrations of the radial motion patterns under various speeds and the task paradigm. (**A**) The expanding (left column) and contracting (right column) optic flow patterns are moving under low (first row), medium (middle row), and high (bottom row) speeds. Dots between panels indicate that multiple speed conditions (>3) are present in the experiments. (**B**) The illustration of the time series of each trial. Participants were asked to perform a speed discrimination task between the reference stimulus and the comparison stimulus. Of note, while Experiments 1 and 2 used different stimulus designs (see **Supplementary Videos 1-4**), we used a similar time series of each trial in both experiments.

#### Procedure

**Figure 1B** depicts the time course of one trial in the experiment. The experiment was conducted with a two-interval forced-choice (2IFC) task. Each trial started with a 0.5-second presentation of the reference stimulus, followed by a 1-second blank screen inter-stimulus interval (ISI) and the other 0.5-second presentation of the comparison stimulus. Participants were later asked to report whether they perceived the comparison stimulus as moving slower or faster than the reference stimulus by pressing a button on the gaming pad. Participants were asked to maintain fixation on the central cross during the task, except during the reporting period. Five speed conditions (1, 4, 16, 64, and 128 °/s) were tested for both expanding and contracting optic flow patterns, referred to as reference stimuli. In each trial of the speed comparison task, a reference stimulus was followed by a comparison stimulus of the same motion pattern but at a different speed. The speeds of the comparison stimulus were selected exponentially (in powers of 2), including three slower, one identical, and three faster than the reference stimulus. The experiment consisted of 35 conditions (5 reference speeds × 7 comparison speeds) for each of the two optic flow patterns (expansion and contraction), resulting in a total of 70 conditions. The session was divided into three blocks, with each condition presented five times per block in a pseudorandomized order. This design resulted in 350 trials per block (70 conditions × 5 repetitions), totaling 1,050 trials (350 trials × 3 blocks), and 15 repetitions per condition across the entire experiment. Each trial lasted approximately 4 seconds and consisted of a 0.5-second reference stimulus, a 1-second ISI, a 0.5-second comparison stimulus, a 0.5-second response period, and a 1.5-second intertrial interval (ITI). The task lasted approximately 70 minutes.

#### Data analysis and quantification

For each participant, we computed the choice proportion (CP), defined as the proportion of trials in which the comparison stimulus was perceived as faster than the reference stimulus. For each reference speed, seven CP values (one for each comparison speed) were used to fit a psychometric function. Stimulus speeds were log_2_-transformed, and the CPs were fitted with a cumulative normal function implemented in the Palamedes Toolbox (Prins & Kingdom, 2018; https://www.palamedestoolbox.org/):

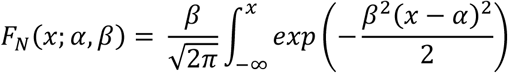

Where 𝐹_𝑁_ is the choice proportion, 𝑥 is the log_2_ transformed comparison speeds, 𝛼 is the threshold, and 𝛽 is the slope of the fitted psychometric curve. The fitting was based on a maximum likelihood criterion evaluated by the following function:

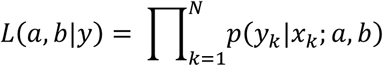

Where 𝐿 is the maximum likelihood, 𝑝 is the probability of observance of response 𝑦 on trial 𝑘 (here the choice proportion) given stimulus intensity 𝑥_𝑘_ (here the comparison speed), and assuming threshold 𝛼 = 𝑎 and slope 𝛽 = 𝑏.

We quantified perceptual sensitivity using the Weber fraction, which describes the smallest relative change in stimulus magnitude that can be discriminated (Gescheider, 2013). For each reference speed, the 50% point of the fitted psychometric function defined the point of subjective equality (PSE). The discrimination threshold was defined as the comparison speed corresponding to 75% choice proportion. The Weber fraction was then computed as:

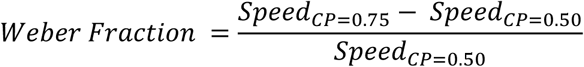

Where 𝑆𝑝𝑒𝑒𝑑_𝐶𝑃=0.75_ is the comparison speed when the choice proportion reaches 75% (discrimination threshold), and 𝑆𝑝𝑒𝑒𝑑_𝐶𝑃=0.50_ is the comparison speed when the choice proportion reaches 50% (PSE). The Weber fraction indicates the minimal percentage change in speed required for reliable discrimination and was used as a standard for evaluating perceptual sensitivity to speed under expansion and contraction conditions.

The distributions of the Weber fractions across 26 participants are illustrated using violin plots generated with the Violinplot MATLAB toolbox (Bechtold, Fletcher, Seamusholden, & Gorur-Shandilya, 2021; https://zenodo.org/records/4559847); the width of each violin represents a kernel density estimate of the data. A boxplot is overlaid on each violin, with the median indicated by a white dot, the box spanning the 25th–75th percentiles, and whiskers extending to 1.5 times the interquartile range. All other boxplots in the Results follow the same setting. Error bars indicate 95% bias-corrected and accelerated (BCa) bootstrap confidence intervals of the median (10,000 resamples).

For each speed condition, we statistically assessed differences in Weber fractions between expansion and contraction using the Wilcoxon signed-rank test. To control for multiple tests performed for speed conditions, statistical significance was evaluated by using the Benjamini–Hochberg false discovery rate (FDR) procedure with *q* = 0.05. At the same time, the uncorrected p-value was reported in Results (presented as *p* in the Results). Effect sizes were quantified using the rank-biserial correlation (presented as *r* in the Results). We additionally report a robust paired Cohen’s d as a standardized measure of effect magnitude (presented as *d* in the Results).

### Results

We examined how accurately observers could discriminate motion speeds under five speed conditions.

**Figure 2A** depicts the results from one participant (top panel, expansion; bottom panel, contraction). The fitted psychometric functions provide a direct visualization of task performance, indicating that this participant can reliably discriminate differences in optic flow speeds, even when the stimulus moves at very slow (1 °/s) or fast (128 °/s) speeds. **Figure 2B** shows the distributions of Weber fractions across flow patterns and speed conditions. For this participant, the Weber fractions are 0.649, 0.419, 0.353, 0.231, and 0.174 for expansion speed conditions of 1, 4, 16, 64, and 128 °/s, and 0.370, 0.319, 0.190, 0.109, and 0.113 for corresponding contraction speed conditions. According to Weber’s law, a smaller Weber fraction indicates higher perceptual sensitivity, as a smaller change in speed is required for a participant to notice the difference. The consistently lower Weber fraction values for contraction than for expansion indicate that this participant had higher perceptual sensitivity to contraction across all speed conditions. Moreover, for both types of optic flow, the Weber fraction decreased monotonically with increasing reference speed, suggesting that the participant’s perceptual sensitivity to speed did not strictly follow Weber’s law.

**Figure 2.**
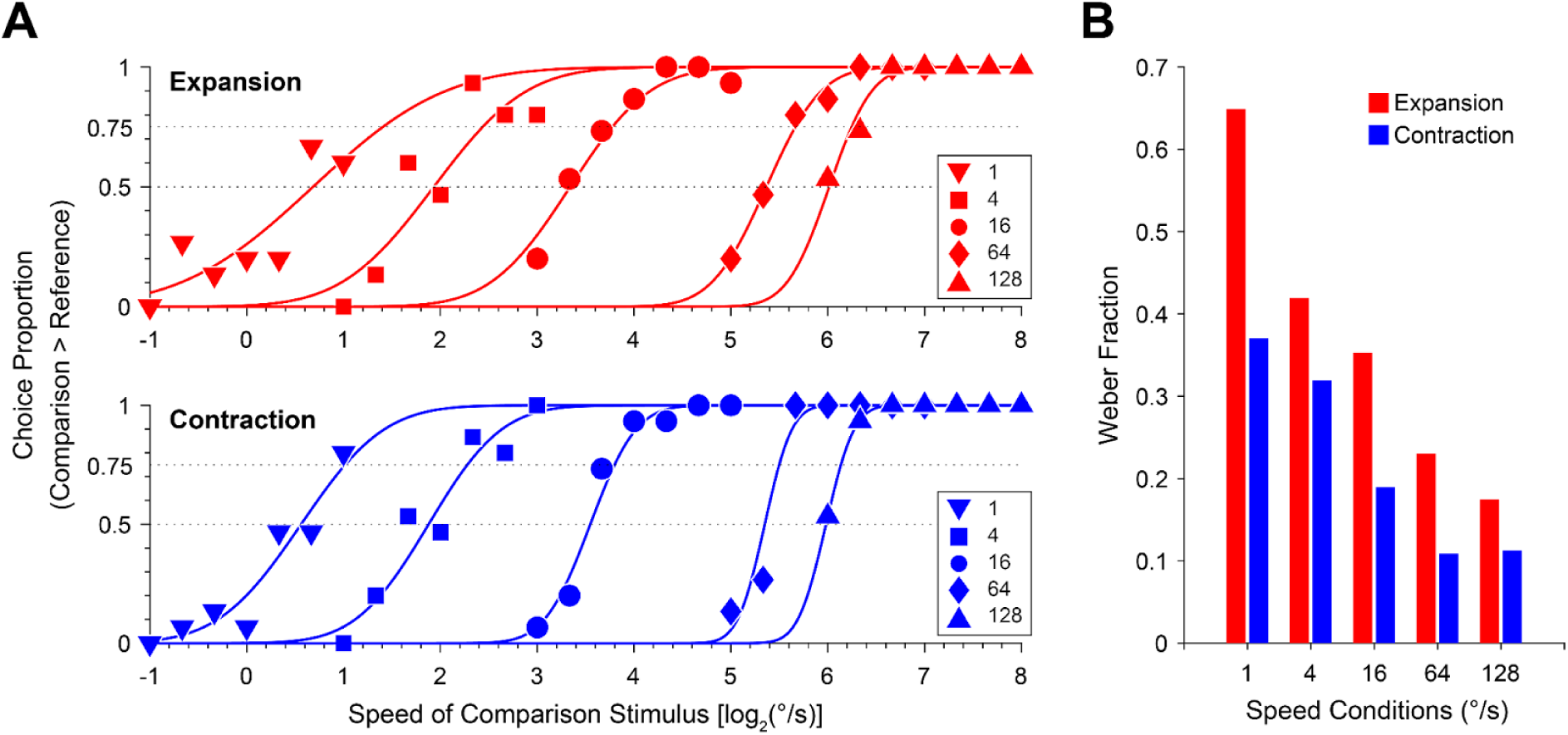
Results of one participant (P12) in Experiment 1. (**A**) The distributions of choice proportions together with the fitted psychometric functions. The dots of different shapes depict the data on choice proportions in each reference speed condition (see legend at the bottom right), whereas the curves represent the fitted psychometric functions (red: expansion conditions; blue: contraction conditions). The horizontal axis shows the log_2_-transformed speeds of the comparison stimulus, whereas the vertical axis shows the choice proportion of reporting that the comparison stimulus moves faster than the reference stimulus. The dashed lines represent the choice proportion thresholds of 50% and 75%, which were used to calculate the Weber fraction (see Method). (**B**) The Weber fractions for expansion (red) and contraction (blue) stimulus at each reference speed. This participant shows lower Weber fractions for contraction than for expansion at all speed conditions, indicating a consistently higher perceptual sensitivity to contraction across speeds.

To determine whether this pattern was representative beyond an individual participant, we next examined the population statistics to assess whether the observed anisotropic perceptual sensitivity is a common perceptual phenomenon.

**Figure 3** depicts the distributions of Weber fractions in all 27 participants. We found significant differences between expansion and contraction at the reference speeds of 4 and 16 °/s. The median Weber fraction of 4 °/s (**Figure 3B**) is 0.411 (95% BCa CI [0.348 0.495]) for expansion and 0.333 (95% BCa CI [0.309 0.446]) for contraction (*p* = 0.007; *r* = 0.598; *d* = 0.470); the median Weber fraction of 16 °/s (**Figure 3C**) is 0.353 (95% BCa CI [0.320 0.441]) for expansion and 0.312 (95% BCa CI [0.251 0.406]) for contraction (*p* = 0.016; *r* = 0.529; *d* = 0.403). In both conditions, the differences remained statistically significant after controlling for the FDR using the Benjamini–Hochberg procedure (*q* = 0.05; five comparisons). The smaller mean Weber fraction for contraction than for expansion indicates that participants are more sensitive to contraction, especially at speeds of 4 or 16 °/s.

**Figure 3.**
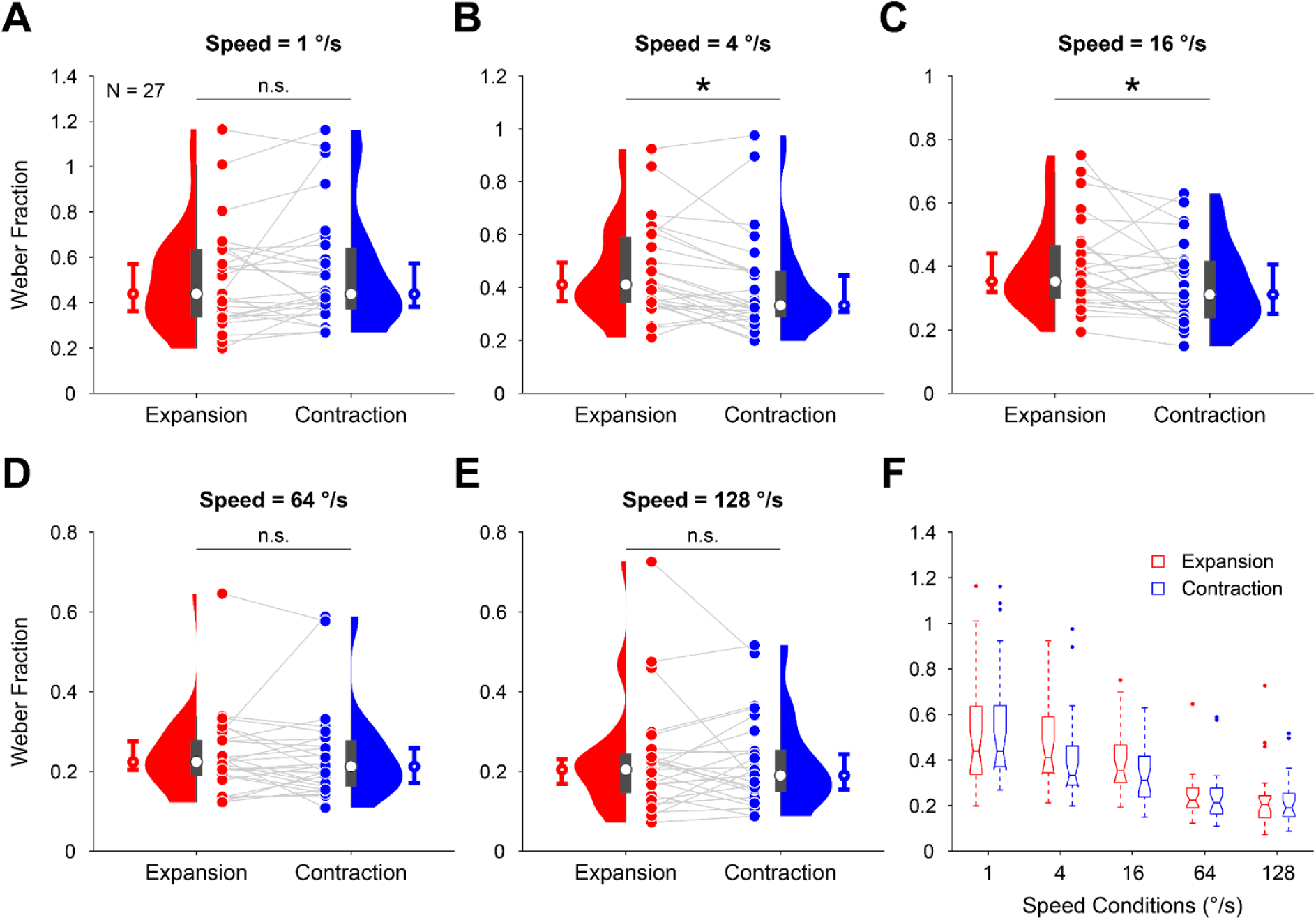
Population results of Weber fraction distributions in Experiment 1 (N = 27). (**A**-**E**) Results of five reference speed conditions. The dots and violin box plots show the distributions of Weber fractions across all participants (see Method for a detailed description of the violin and box plots). The white dots representing the sample median are overlaid on the box plots (gray) that depict the 25th-75th percentiles (thick boxes) and 1.5 times the interquartile range (thin lines). Error bars beside the violin plots depict the 95% BCa bootstrap confidence intervals of the median (open circles overlaid on the error bars). Red represents the results of expansion conditions, and blue represents the results of contraction conditions. Data points from the same participant are connected by light gray lines. Despite comparable mean Weber fractions for expansion and contraction at low and fast speeds, perceptual sensitivities to contraction are significantly higher than to expansion at reference speeds of 4 and 16 °/s. Asterisks indicate statistically significant differences that survived correction for multiple comparisons using the Benjamini–Hochberg FDR (*q* = 0.05). (**F**) The summaries of the Weber fraction distributions of all reference speed conditions, the horizontal lines in the box plot represent the sample median (see Methods for details of the box plot).

On the other hand, we found comparable perceptual sensitivities to expansion and contraction at reference speeds of 1 °/s, 64 °/s, and 128 °/s. Under the condition of 1 °/s (**Figure 3A**), the median Weber fraction is 0.439 (95% BCa CI [0.363 0.570]) for expansion, and 0.438 (95% BCa CI [0.382 0.573]) for contraction (*p* = 0.313; *r* = -0.222; *d* = -0.067); under the condition of 64 °/s (**Figure 3D**), the median Weber fraction is 0.224 (95% BCa CI [0.203 0.276]) for expansion, and 0.212 (95% BCa CI [0.171 0.258]) for contraction (*p* = 0.597; *r* = 0.116; *d* = 0.132); under the condition of 128 °/s (**Figure 3E**), the median Weber fraction is 0.205 (95% BCa CI [0.168 0.231]) for expansion, and 0.190 (95% BCa CI [0.155 0.243]) for contraction (*p* = 0.848; *r* = -0.042; *d* = -0.026).

These results show that there are anisotropic perceptual sensitivities between expansion and contraction. Still, these differences occur only at medium speed (4 and 16 °/s), which are approximately the egomotion speeds for walking and driving, but not at slow (1 °/s) or fast (64 or 128 °/s) speeds.

## Experiment 2

In Experiment 1, we found anisotropies in perceptual sensitivity between planar expansion and contraction radial flow patterns, with speeds strictly controlled and uniform across visual eccentricity. In the next step, we designed visual stimuli that simulate the 3D optic flow patterns projected onto the screen and tested whether the sensitivity asymmetry found in Experiment 1 was also generalizable to 3D optic flow patterns using the same task paradigm.

### Method

#### Participants

Thirty healthy human adults (19 females and 11 males, aged 33.30 ± 1.74 S.E. years, ranging from 20 to 50 years old) participated in the experiments. All other procedures, such as participant recruitment and screening, informed consent, and ethics committee approval, are the same as those in Experiment 1.

#### Apparatus

Motion stimuli were generated by a gaming desktop computer (Dell Alienware Aurora R16, Dell Inc., Round Rock, TX, USA) running a Linux operating system (Ubuntu 22.04 LTS). Visual stimuli were presented on an LCD monitor (Alienware AW2524HF, Dell Inc., Round Rock, TX, USA) with a refresh rate of 480 Hz and a spatial resolution of 1920 × 1080 pixels. The monitor is 54.4 cm wide and 30.3 cm high. Screen luminance was measured using a colorimeter (CS-150, Konica Minolta Inc., Japan). The minimum (black level) and maximum (white level) luminances are approximately 0.4 and 450 cd/m2, respectively. The distance between the eyes and the projection screen is 65 cm, resulting in a visual angle of approximately 45.52° (width) × 26.36° (height). We note that we used a different apparatus setup in Experiment 2 because the PROPixx projector was unavailable at the time. All experiments were conducted in a dimly dark room. All other procedures are the same as those in Experiment 1.

#### Visual stimuli

In Experiment 2, the random-dot field presented on the screen simulated 3D optic flow in a realistic environment (**Figure 4**), a widely used stimulus design in the literature (Cuturi & MacNeilage, 2014; Danz, Angelaki, & DeAngelis, 2020; Fujimoto & Ashida, 2020). Each dot has a 3D diameter of approximately 115 mm, a dot density of 10^-9.6^ dots/mm^3^, and the speed in 3D space is defined in meters per second (m/s). In the 3D space, all the above parameters were consistent between dots and across the time course of stimulus presentation. The position of each dot on the screen was simulated as a projection of its 3D location along the viewing trajectory. The size of each dot also underwent the same simulation process. As a result, the projected dots on the screen had different moving speeds and sizes across the visual eccentricities, despite their constant speed and size in the 3D space (**Figure 4A**). In detail, the speeds accelerated and the sizes magnified from the fovea to the periphery in a nonlinear manner (**Supplementary video 3**; see **Supplementary video 4** for an example of contracting optic flow). The physical radial distances from the fixation center range from 0.5 to 5 m. The viewing depth ranges from the screen to 100 m beyond it. So, the entire visual stimulus pattern was structured as a star field moving in a cylinder of a given thickness (**Figure 4B**), and a similar structure was also used in the previous study (Fujimoto & Ashida, 2020). Each dot traveled through the entire viewing depth before being removed and reinitialized at the start point, which can be the maximum viewing depth for an expanding flow or the screen for a contracting flow. Same as Experiment 1, a white fixation cross was presented at the center of the screen during the task.

**Figure 4.**
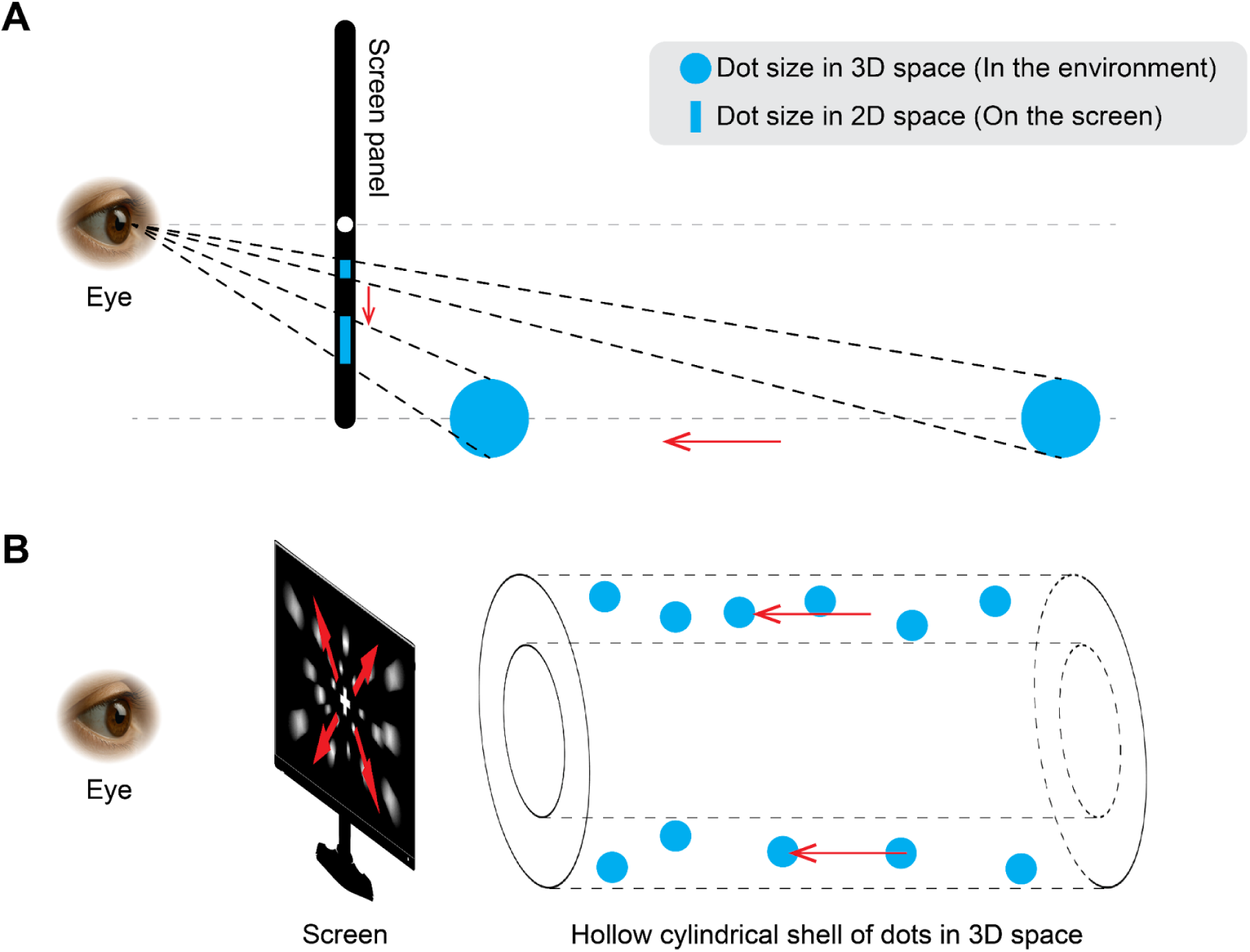
Schematic illustration of stimulus design in Experiment 2, with expanding optic flow as an example. (**A**) The simulation of 3D optic flow patterns on the 2D screen. In the expansion condition, each dot (blue) maintains a constant size in 3D as it approaches the screen (the horizontal red arrow indicates the motion direction). To simulate the stimulus on a 2D screen, dots at different depths were projected onto the screen (blue sticks) based on the participant’s viewpoint. A 3D dot approaching the screen appears as a 2D dot moving away (vertical red arrow) from the fixation point (white dot), with size and speed varying by eccentricity. ( **B**) The visual stimulus consisted of uniform-sized dots moving at a constant speed within a 3D tunnel-shaped cylinder. By applying the method described in (A), dots were rendered with varying sizes and speeds on the screen, simulating a 3D optic flow. We used expanding and contracting 3D optic flow with this design in Experiment 2.

#### Procedure

As in Experiment 1, Experiment 2 used a 2IFC task with a similar time-course design. Instead of fixed comparison speed steps, we used the Psi adaptive staircase method, which is robust and efficient for estimating both thresholds and slopes (Kontsevich & Tyler, 1999), to automatically select the comparison speed from a predefined speed range. Four 3D-space reference speed conditions (0.5, 1.5, 20, and 100 m/s) were tested for both expansion and contraction. There are 30 trials for each reference speed condition. The selection of the comparison speed is determined by the Psi staircase procedure and based on the prior task responses. As a result, Experiment 2 consisted of 240 trials (4 reference speeds × 2 optic flow patterns × 30 trials). Each trial lasted approximately 6.5 seconds and consisted of a 1-second reference stimulus, a 2-second ISI, a 1-second comparison stimulus, a 0.5-second response period, and a 2-second ITI. The total task duration was approximately 26 minutes.

#### Data analysis and quantification

The data analysis was conducted in the same way as in Experiment 1.

### Results

The 3D-space reference speeds in Experiment 2 were purposely set to be 0.5, 1.5, 20, and 100 m/s, where 0.5 m/s is smaller than most of the egomotion compatible speeds, 1.5 m/s (5.4 km/h) and 20 m/s (72 km/h) simulate the approximate speeds of walking and driving, respectively, and 100 m/s (360 km/h) is faster than a bullet train. These speeds provide an efficient range of stimulus parameters for testing whether anisotropic perceptual sensitivity between expansion and contraction exists in well-experienced daily-life body situations and whether the same tendency persists in other, less-experienced speed conditions.

Similar to Experiment 1, the results for one participant show clean, well-defined psychometric function curves across all 3D-simulated expansion and contraction speed conditions (**Figure 5A**). Markedly, the perceptual sensitivities of contraction are higher than expansion (**Figure 5B**) at the reference speeds of 1.5 m/s (Weber fraction is 0.269 for expansion and 0.226 for contraction) and 20 m/s (Weber fraction is 0.131 for expansion and 0.064 for contraction), which simulate the egomotion condition of walking and driving. In contrast, the tendencies are opposite under conditions of 0.5 m/s (Weber fraction is 0.202 for expansion and 0.266 for contraction) and 100 m/s (Weber fraction is 0.081 for expansion and 0.109 for contraction).

**Figure 5.**
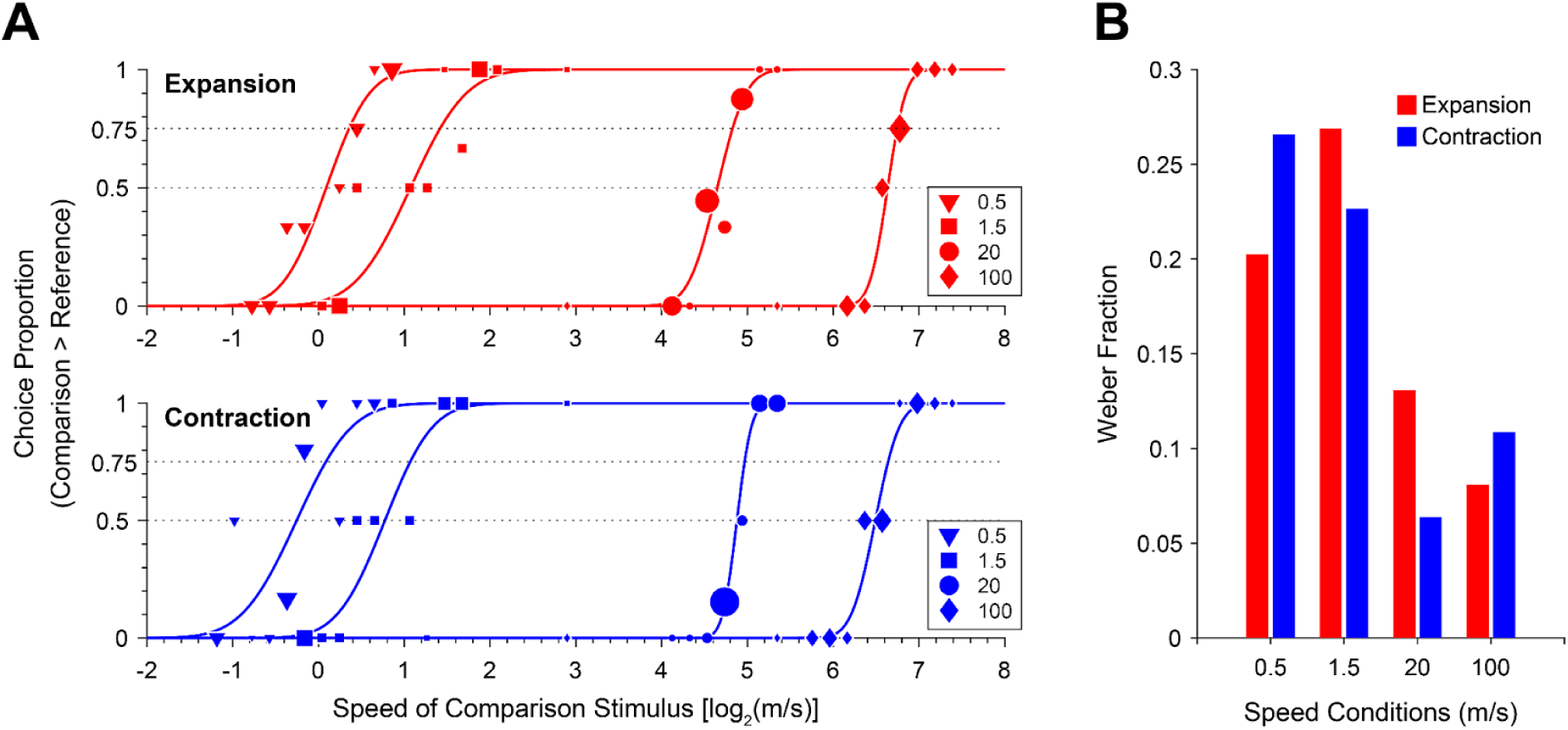
Results of one participant (P13) in Experiment 2. (**A**) The distributions of choice proportions together with the fitted psychometric functions. The size of each data point reflects the number of repetitions for the corresponding comparison speed in the adaptive staircase task. Larger sizes indicate more repetitions. Other conventions are identical to those in Figure 2A. (**B**) The Weber fractions for expansion (red) and contraction (blue) stimulus at each reference speed. This participant shows higher perceptual sensitivities to contraction at reference speeds of 1.5 and 20 m/s, but opposite tendencies at 0.5 and 100 m/s. Other conventions are identical to those in Figure 2B.

In the population data of 30 participants, we found a significantly higher perceptual sensitivity to contracting than expanding speed under walking speed (1.5 m/s) condition (**Figure 6B**, median Weber fraction of expansion: 0.573, 95% BCa CI [0.414 0.724]; median Weber fraction of contraction: 0.426, 95% BCa CI [0.367 0.515]; *p* = 0.010, *r* = 0.540, *d* = 0.539). This difference remained significant after controlling for the FDR using the Benjamini–Hochberg procedure (*q* = 0.05; four comparisons).

**Figure 6.**
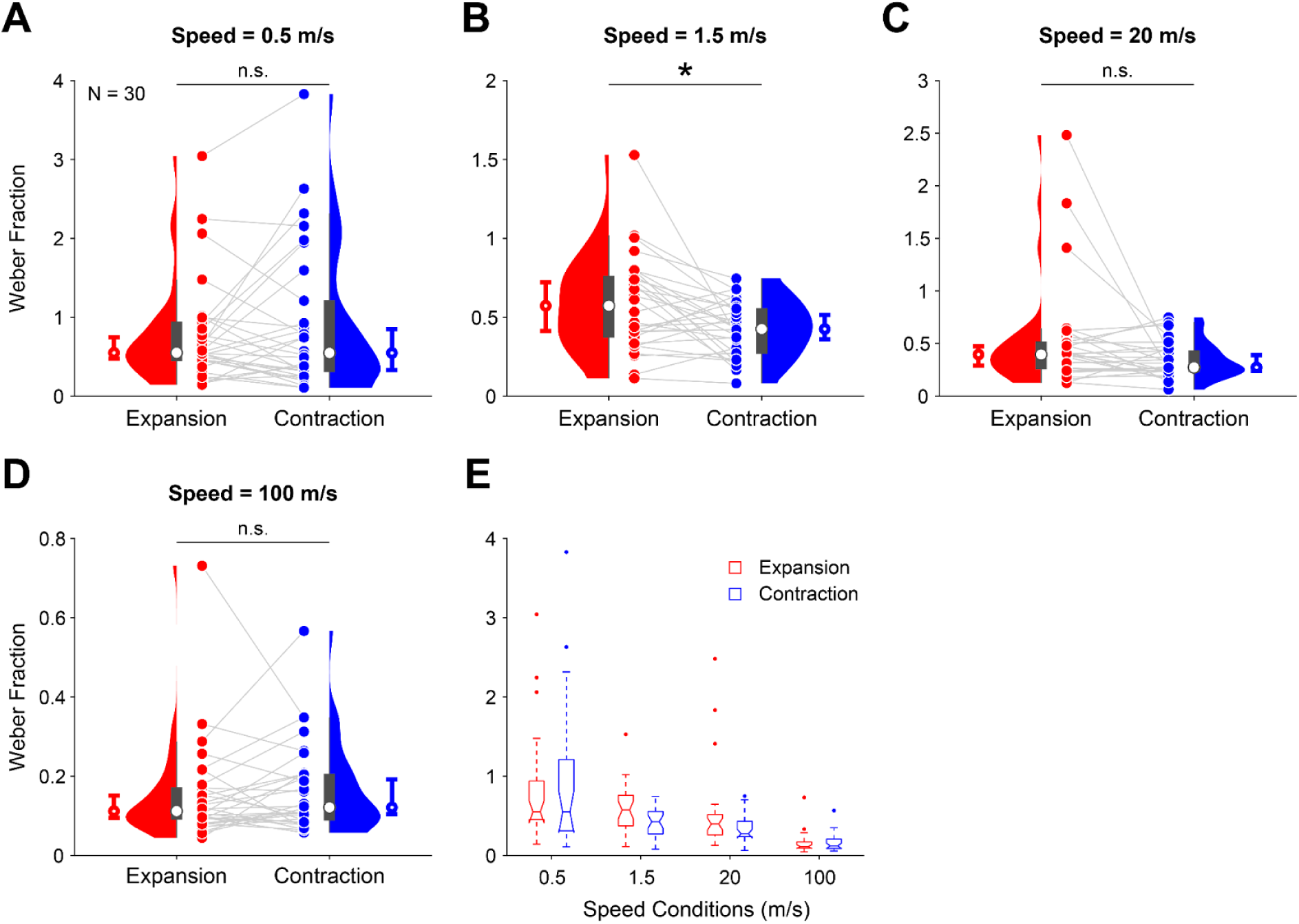
Population results of Weber fraction distributions in Experiment 2 (N = 30). (**A**-**D**) Results of four reference speed conditions. Perceptual sensitivities to contraction are significantly higher than to expansion at the walking speed of 1.5 m/s, but not at other speeds. The asterisk indicates a statistically significant difference that survived correction for multiple comparisons using the Benjamini–Hochberg FDR (*q* = 0.05). (**E**) The summaries of the Weber fraction distributions of all reference speed conditions. The other conventions are identical to those used in Figure 3.

In contrast, no significant anisotropy was found in speed conditions of 0.5 m/s (**Figure 6A**, median Weber fraction of expansion: 0.551, 95% BCa CI [0.478 0.749]; median Weber fraction of contraction: 0.549, 95% BCa CI [0.336 0.852]; *p* = 0.673, *r* = -0.088; *d* = -0.175), 20 m/s (**Figure 6C**, median Weber fraction of expansion: 0.397, 95% BCa CI [0.292 0.480]; median Weber fraction of contraction: 0.275, 95% BCa CI [0.241 0.392]; *p* = 0.178, *r* = 0.282; *d* = 0.390), and 100 m/s (**Figure 6D**, median Weber fraction of expansion: 0.112, 95% BCa CI [0.095 0.152]; median Weber fraction of contraction: 0.121, 95% BCa CI [0.104 0.192]; *p* = 0.319, *r* = -0.209; *d* = -0.224). Additionally, as with planar radial flow patterns (Experiment 1), the Weber fractions of 3D simulated optic flow patterns also decreased from low to high speeds (**Figure 6E**).

In summary, Experiment 2 demonstrated that perceptual sensitivity to contraction was significantly higher than that to expansion when stimuli simulated 3D optic flows at a speed corresponding to the walking speed (1.5 m/s). Despite differences in stimulus design and experimental setup between the two experiments, both showed higher sensitivity to contraction at medium speed. This convergence indicates that anisotropic processing of expanding versus contracting optic flows is robust across display formats and may reflect a general visual mechanism tuned to ecologically relevant patterns of self-motion, such as walking.

## Discussion

This study investigated the speed-discrimination abilities of human participants for expanding and contracting radial optic flow, ranging from 1 to 128 °/s for 2D radial flow patterns (Experiment 1) and from 0.5 to 100 m/s for 3D optic-flow patterns (Experiment 2). We used the Weber fraction as a criterion to evaluate the perceptual sensitivities of the two flow patterns under various speeds. Results showed that perceptual sensitivities to contraction are higher than those to expansion at speeds of 4 and 16 °/s under 2D conditions and 1.5 m/s under 3D conditions. In contrast, this between-pattern anisotropy was not significant under other lower and higher reference speed conditions. Below, we discuss the significance of these findings from a psychophysical and ecological perspective.

### The controversies in perceptual sensitivities between psychophysical studies

As mentioned in the Introduction, different studies yielded conflicting conclusions regarding perceptual sensitivity to expansion and contraction. Specifically, while some works reported greater perceptual sensitivity to contraction, which is consistent with our results (Edwards & Badcock, 1993; Edwards & Ibbotson, 2007; Geesaman & Qian, 1998), others found opposite results (Clifford et al., 1999; Lopez-Moliner, 2005) or no clear difference (Beardsley & Vaina, 2005; Morrone et al., 1999). Here, we discuss the potential reasons for this apparent inconsistency across psychophysical studies from three perspectives.

One key source of inconsistency likely lies in the different stimulus design. Several studies employed large-field random-dot optic flow stimuli (Clifford et al., 1999; Edwards & Badcock, 1993; Edwards & Ibbotson, 2007; Geesaman & Qian, 1998), which generate genuinely global expansion or contraction patterns resembling those produced during self-motion. In contrast, Lopez-Moliner (2005) used an expanding or contracting square, which is spatially restricted and conveys less self-motion information than optic flow, which provides more ecologically plausible information. Thus, variation in stimulus type offers a plausible explanation for why studies sometimes report conflicting conclusions about whether perceptual sensitivity is higher for expansion or contraction.

A second source of inconsistency across studies concerns the task paradigms used. Edwards & Badcock (1993) and Edwards & Ibbotson (2007) estimated sensitivity by determining the number of signal dots required to achieve 75% correct performance. Clifford et al. (1999) assessed speed discrimination by comparing multiple reference optic flow patterns against a single comparison pattern. In contrast, Lopez-Moliner (2005) used a detection paradigm and measured reaction times for expansion, contraction, and translational motion. These approaches differ not only in the stimulus features used to infer sensitivity (e.g., signal-dot proportion vs. speed) but also in the behavioral measures involved (discrimination thresholds vs. response latencies). Because these metrics tap distinct perceptual and decision processes, differences in task design may also contribute to conflicting conclusions about relative sensitivity to expansion and contraction.

A third factor concerns the limited range of stimulus parameters used in previous work, particularly speed. Although many studies have compared perceptual sensitivity to expansion and contraction, only a few have examined how this anisotropy varies systematically across different speeds (e.g., Edwards & Ibbotson, 2007), and even then, the tested range was relatively narrow (2.4–28.8°/s). Only two studies explored perceptual sensitivity across a broader speed range (De Bruyn & Orban, 1988; Orban, de Wolf, & Maes, 1984), but both used translational motion rather than radial optic flow, making their findings difficult to generalize to expansion and contraction. Our results, showing significant anisotropy, were observed only in a specific speed range, providing a plausible explanation for why some studies have not observed clear differences between expansion and contraction. For example, Beardsley & Vaina (2005) tested speeds of only 8.4°/s and 30°/s, and Morrone et al. (1999) used stimuli with a speed gradient peaking at 4°/s (although their study did not primarily aim to compare radial flow types). Studies sampling only a narrow or ecologically unrepresentative speed range may therefore miss speed-dependent differences in radial optic flow processing.

To our knowledge, this is the first study to systematically examine perceptual sensitivity to radial optic flow across such a broad range of speeds using a large-field random-dot stimulus, which is well suited for probing global motion mechanisms by overcoming the third factor discussed above. By leveraging a state-of-the-art display system with a 1440-Hz projection rate, we were able to present motion patterns that approximate natural-scene computable speeds, including both extremely slow and exceptionally fast optic flows. This methodological advantage allowed us to characterize how perceptual sensitivity evolves across the full speed continuum and to demonstrate that anisotropic sensitivity between expansion and contraction is present, but primarily within the medium-speed range.

More broadly, the current findings highlight the flexibility and context-dependence of the visual system when processing radial motion. Although resolving the source of all inconsistencies is not practical for a single study, future studies using different stimulus types, task structures, and speed ranges can further probe distinct motion-processing mechanisms, thereby helping explain inconsistencies across previous studies.

### The anisotropic perceptual sensitivity was generally observed in both 2D and 3D optic flow patterns

Previous psychophysical work has shown that perceptual sensitivity to radial optic flow patterns is anisotropic for large-field 2D motion stimuli (Clifford et al., 1999; Edwards & Ibbotson, 2007; Shirai, Kanazawa, & Yamaguchi, 2006). Additionally, some functional MRI (fMRI) studies found that motion-selective cortical areas respond differently to optic flow with and without stereoscopic speed cues (Cardin & Smith, 2011; Uesaki & Ashida, 2015), suggesting that psychophysical findings from 2D optic flow may not generalize to 3D optic flow.

However, to our knowledge, no previous study has directly compared anisotropic speed discrimination for expanding and contracting optic flows in both 2D and 3D conditions within a single experimental framework. The present results, therefore, extend the existing literature by linking directional anisotropies in radial flow to differences between 2D and 3D optic flow processing. Despite differences in stimulus patterns, higher perceptual sensitivity to contraction relative to expansion was consistently observed at medium (walking) speed in both 2D and 3D conditions. Together, these findings demonstrate that anisotropic perceptual sensitivity to radial optic flow is not restricted to simplified 2D displays but extends to naturalistic 3D flow fields resembling those experienced during self-motion.

Of note, results for 3D optic flow (Figure 6) exhibited greater inter-participant variability and lower sensitivity than those for 2D optic flow (Figure 3). This may be explained by higher task difficulty for discriminating 3D optic flow speed, given that dot speed and size varied systematically across visual eccentricity.

### The ecological significance of perceptual anisotropy

In our study, anisotropic perceptual sensitivity was evident at egomotion speeds for walking and driving (only for walking speed with 3D patterns), with the most substantial effect observed at walking speed. The absence of such differences at much lower or higher speeds is unsurprising, as such speeds are less common in everyday visual experience and are likely to have limited biological relevance. However, it remains unclear why perceptual sensitivity to contraction exceeds that to expansion at walking speed, given that expanding optic flow is generally more ecologically significant. The anisotropic visual sensitivities are heavily derived from the anisotropic environmental stimulations (Karim & Kojima, 2010). For humans, the most frequently experienced self-motion is walking forward, which generates an expanding flow pattern on the retina. Therefore, evolutionary and developmental factors should lead to greater perceptual sensitivity to expansion than to contraction.

One possible explanation for the higher sensitivity of contraction relies on its specific visual functionality. One typical case of normal visual anisotropy is the oblique effect, which describes the fact that our visual system is better at discriminating orientations along the vertical and horizontal cardinal axes than along oblique axes (Appelle, 1972; Furmanski & Engel, 2000). This perceptual anisotropy is attributed to the adaptive properties of the visual system, which respond to the environment and provide rich visual information along cardinal axes (Annis & Frost, 1973). In this case, perceptual sensitivity aligned with the visual experience, and one factor that could lead to this alignment is that oblique orientations are of less biological importance. In the case of egomotion, contracting optic flow usually reflects important self-body conditions and may be used to control body balance and to alert to dangerous body situations. Notably, the speeds of 4, 16 °/s and 1.5 m/s approximate typical self-motion scenarios such as walking/running and driving, which are the most experienced body movements in daily life, instead of the expanding flow which is generated in those normal conditions, contracting flow can be induced by abnormal body postures such as falling onto our back when we are walking and facing backward when we are driving. Under those conditions, contracting optic flow is typically considered a sign of danger (**Figure 7**). Increasing the perceptual sensitivity of contraction in our visual system can alert us to potentially hazardous body conditions and aid in rapid reactions to maintain body balance and normal posture. In contrast, the absence of anisotropy at higher speeds implies that this advantage may be limited to ecological regimes in which humans usually rely more heavily on visual self-motion cues.

**Figure 7.**
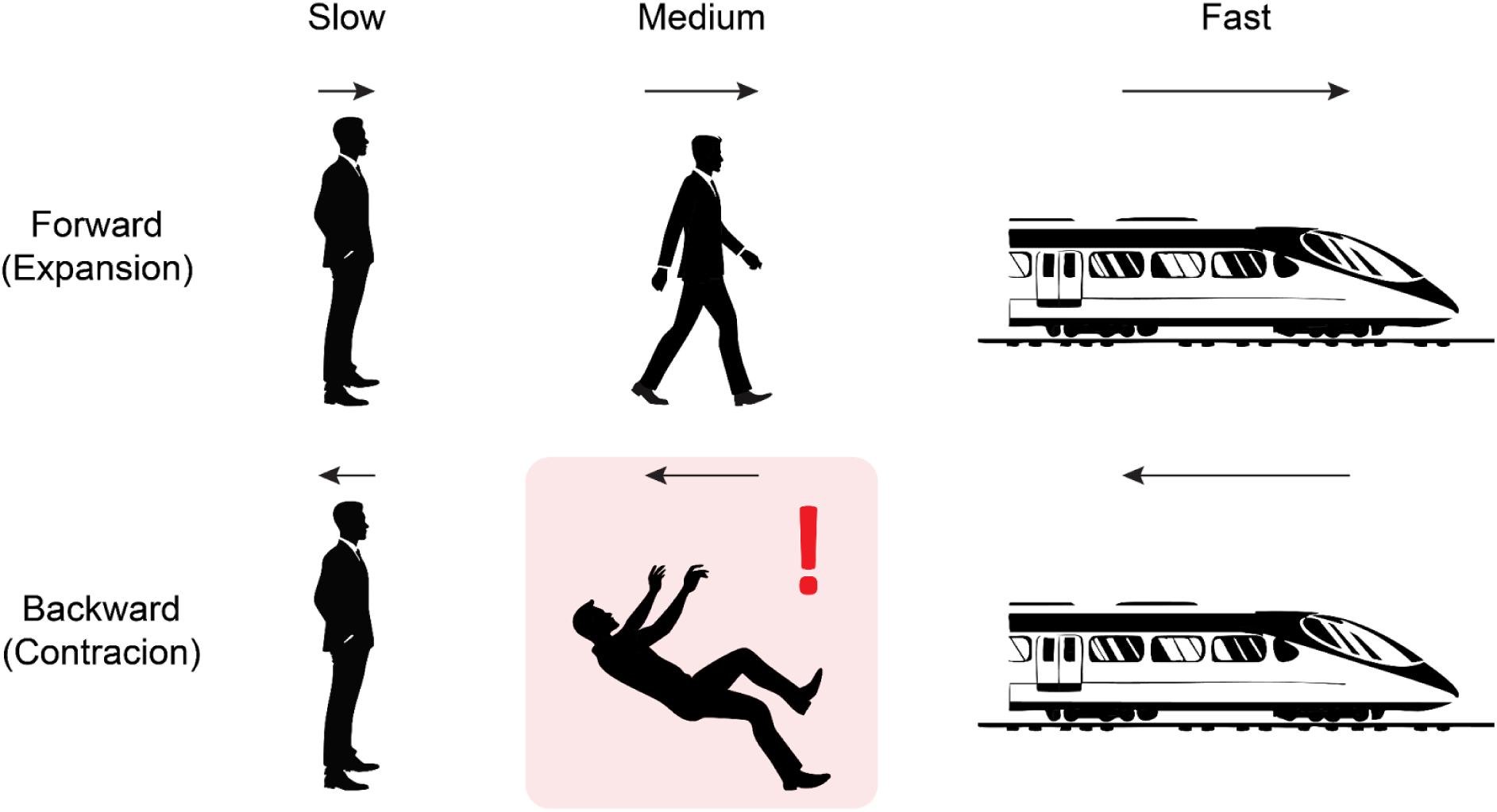
Schematic illustration of optic flow associated with different egomotion states. Forward locomotion produces expanding optic flow, whereas backward motion produces contracting optic flow, with flow speed varying systematically from slow to fast. Among these conditions, backward falls during locomotion elicit medium-speed contractions that typically signal imminent danger and require rapid visuomotor responses. The ecological importance of detecting such events may help explain the heightened perceptual sensitivity to contracting optic flow.

Notably, prior studies of vection, the subjective illusion of self-motion induced by optic flow, have reported that contraction elicits stronger effects than expansion (Bubka, Bonato, & Palmisano, 2008; Fujimoto & Ashida, 2019). These findings are consistent with our results regarding the ecological interpretation of anisotropy, while testing a direct relationship between speed sensitivity and subjective perception of vection is a topic for future investigation.

### The remaining questions and future perspectives

Perceptual sensitivity to radial optic flow was compared in this study, but egomotion-induced flow patterns are not limited to expansion and contraction. Body movement conditions, such as turning around a corner while walking forward, usually induce rotary optic flow patterns. Studies have reported that the perceptual speed, or say perceptual sensitivity, of rotary optic flow is lower than that of radial optic flow (Beardsley & Vaina, 2005; Clifford et al., 1999; Geesaman & Qian, 1998; Morrone et al., 1999). However, it still needs further exploration to determine whether this perceptual anisotropy persists across different speed ranges and whether it exists between clockwise and counterclockwise optic flow patterns.

Electrophysiological studies have found an overrepresentation of expansion-sensitive neurons relative to contraction-sensitive neurons in macaque area MSTd, which is highly selective for optic flow patterns (Duffy & Wurtz, 1991; Graziano, Andersen, & Snowden, 1994; Tanaka et al., 1989). These findings apparently conflict with our psychophysical data, which show higher sensitivity to contraction. However, a larger number of neurons for a specific type of stimulus does not always yield better behavioral performance, as the relationship between population coding and behavioral sensitivity depends on multiple factors, including tuning width (Kumano & Uka, 2023) and correlations across neurons (Averbeck, Latham, & Pouget, 2006; Zohary, Shadlen, & Newsome, 1994). Understanding the underlying relationship between neuronal and perceptual mechanisms is an essential topic for future studies.

Compared with macaque electrophysiology, the understanding of mechanisms for speed encoding in the human brain remains limited, although previous studies have investigated this topic by using magnetoencephalography (MEG) (Amano, Kimura, Nishida, Takeda, & Gomi, 2009; Kawakami et al., 2002) and fMRI (Lingnau, Ashida, Wall, & Smith, 2009; Nishimoto et al., 2011; Takemura, Ashida, Amano, Kitaoka, & Murakami, 2012; Vintch & Gardner, 2014). However, since none of these studies used optic flow as a visual stimulus, the exact mechanisms underlying the speed representation of optic flow in humans remain unknown. Extension of this study, using MEG or fMRI, can provide further insights into how anisotropies in perceptual sensitivity are associated with responses in motion-selective areas in humans.

## Conclusion

In this study, we demonstrated anisotropic perceptual sensitivity to different radial optic flow patterns across a specific speed range, in both 2D and 3D optic flow. Notably, this anisotropy is insignificant at both lower and higher speed ranges. It emerges most clearly at medium speeds most frequently experienced in everyday self-motion, such as walking or driving, suggesting that the visual system may be adaptively tuned to ecologically relevant motion conditions.

## Acknowledgments

We thank Kumiko Kobayashi and Yuri Morii for their assistance in participant recruitment and data collection.

## Data and code availability

The data and code for replicating figures and statistical results are available in the public repository (https://github.com/OkazakiTakemuraLab/PerceptualAnisotropy).

## Supplementary materials

**Supplementary Video 1**. This video illustrates the 2D planar expanding optic flow stimulus employed in Experiment 1. The radial motion expands at a uniform speed of 4 °/s.

**Supplementary Video 2**. This video illustrates the 2D planar contracting optic flow stimulus employed in Experiment 1. The radial motion contracts at a uniform speed of 4 °/s.

**Supplementary Video 3**. This video illustrates the expanding optic flow stimulus employed in Experiment 2, designed to emulate the expanding optic flow experienced in natural environments. The simulated motion speed is 1.5 m/s.

**Supplementary Video 4**. This video illustrates the contracting optic flow stimulus employed in Experiment 2, designed to emulate the contracting optic flow experienced in natural environments. The simulated motion speed is 1.5 m/s.

